# Event-Induced Modulation of Aperiodic Background EEG: Attention-Dependent and Age-Related Shifts in E:I balance, and Their Consequences for Behavior

**DOI:** 10.1101/2023.04.14.536436

**Authors:** Patrycja Kałamała, Máté Gyurkovics, Daniel C. Bowie, Grace M. Clements, Kathy A. Low, Florin Dolcos, Monica Fabiani, Gabriele Gratton

## Abstract

The broadband shape of the EEG spectrum, summarized using a 1/*f^x^* function, is thought to reflect the balance between excitation and inhibition in cortical regions (E:I balance). This balance is an important characteristic of neural circuits and could inform studies of aging, as older adults show a relative inhibitory activity deficit. Thus far, no studies have leveraged the event-related temporal dynamics of 1/*f*^x^ activity to better understand the phases of information processing, especially in the context of aging. Here, for the first time, we examined variations of this activity during the foreperiod of a cued flanker task in younger (YA) and older adults (OA), with picture cues varying in task relevance, relative novelty, and valence. We report a biphasic change in the spectral exponent (corresponding to negative slopes in log-log space) after cue presentation, independent of cue-elicited ERPs, with an initial period of increased negativity (indicating cortical inhibition, similar in YA and OA) followed by decreased negativity (indicating cortical excitation, especially in OA). The decrease in the exponent negativity was associated with lower performance and greater congruency costs in the flanker task. Finally, more novel cues reduced the shift towards excitation in OA, partly restoring their E:I balance, and diminishing congruency costs. These findings demonstrate that the broadband shape of the EEG spectrum varies dynamically in a manner that is predictive of subsequent behavior. They also expand our understanding of how neural communication shapes cognition in YA and OA and have implications for neuroscientific models of cognitive processing and age-related cognitive decline.

## 1. Introduction

The brain constantly exhibits a repertoire of complex dynamics related to behavior in health and disease. In the electrophysiological power spectrum, brain dynamics are expressed in the form of oscillatory/periodic voltage fluctuations, emerging against non-oscillatory/aperiodic background activity. Despite accounting for a substantial portion of the neural signal, the aperiodic component has, until recently, received limited attention in cognitive neuroscience, often being considered “noise” devoid of any functional significance. Recent theoretical and methodological advances, however, have begun to provide evidence in support of the functional relevance of the aperiodic component in explaining brain dynamics and human behavior (Donoghue et al., 2020; Gyurkovics et al., 2022; Voytek et al., 2015; Waschke et al., 2021). A significant breakthrough in this research is the observation of a reduction in the exponent of the aperiodic activity (i.e., a flatter spectrum) in older adults, consistent with the idea of increasing neural noise in aging (Cremer & Zeef, 1987; Salthouse, 2010; Salthouse & Lichty, 1985; Voytek & Knight, 2015), and suggesting an age-related shift in the balance between excitation and inhibition (E:I balance, Gao et al., 2017). In this article, we underscore the rich exogenous and endogenous features of scalp-recorded aperiodic neural activity and show, for the first time, evidence for its dynamic alternations over time. Crucially, these dynamics differ between younger and older adults and correlate with behavioral performance.

Aperiodic neural activity (also called 1/*f* noise) is characterized by a progressive decrease in power across increasing frequencies, which follows a 1/*f^x^* function, where *f* denotes frequency, and *x* is a spectral exponent that can be estimated from the steepness of the power decay in log-log space. Because the aperiodic component follows an inverse power function, its parameters (exponent and offset) are best characterized by using log-log power spectra, where they can be estimated from the negative slope and the intercept of the background spectrum (once periodic components are subtracted). In this article, spectral exponents *x* were characterized in log-log space following the equation of log(1/*f*^x^) = *-x*\*log(*f*). Therefore, a more negative exponent value is associated with a steeper slope, reflecting a shift in power from high to low frequencies, and a less negative value, indicating a shift from low to high frequencies. These exponent changes can also be described as rotations of the log-log power spectrum that are either clockwise (more negative exponent, steeper spectrum) or counterclockwise (less negative exponent, flatter spectrum).

Recent *in silico* modeling, supported by *in vivo* experiments (Ahmad et al., 2022; Cohen & Maunsell, 2011; Gao et al., 2017; Harris & Thiele, 2011; Kanashiro et al., 2017) has shown that the spectral exponent can provide information about the balance between excitatory and inhibitory synaptic circuits (E:I balance), with more or less negative exponents reflecting increased inhibition or excitation, respectively. The spectral exponent *x* can also be interpreted as an index of the degree of synchronization of neural networks during their firing. This suggests that a less negative exponent (i.e., *relatively* greater power at high frequencies) reflects more asynchronous (i.e., noisier) neural communication (Chini et al., 2022; B. J. He, 2014; W. He et al., 2019; Voytek & Knight, 2015). The presented interpretations complement each other and together provide a more complete explanation of the spectral exponent function.

Within these synergistic frameworks, accumulating evidence shows that the spectral exponent obtained from noninvasive EEG recordings can reliably and validly reflect the functional properties of aperiodic neural activity across broad regions of the human cortex (Donoghue et al., 2020; Waschke et al., 2021; Zhang et al., 2023). Consistent with a neural noise hypothesis of aging (Cremer & Zeef, 1987; Salthouse & Lichty, 1985; Voytek & Knight, 2015), several studies have shown a reduced (less negative) exponent for older compared to younger adults, indicating disrupted (noisier) neural communication with advancing age (Clements et al., 2021; W. He et al., 2019; Hill et al., 2022; Merkin et al., 2023; Ostlund et al., 2022). Drawing on the E:I balance framework, the reduced exponent for older adults suggests *an age-related counterclockwise spectral rotation*, signifying an increasing E:I ratio in the aging brain, possibly reflecting a deficit of inhibitory circuits in older adults (see also Gordon et al., 2014).

There is also emerging evidence suggesting that individual differences in the spectral exponent may contribute to age-related cognitive decline, with a reduced exponent associated with poorer outcomes across the adult lifespan (e.g., Dave et al., 2018; Tran et al., 2020; Voytek et al., 2015). This evidence suggests that the increase in neural noise observed in aging – indexed by a decreasing exponent and an increasing E:I ratio – may hamper older adults’ ability to process information. However, the mechanisms behind these phenomena remain elusive, as aperiodic activity is typically derived from the EEG signal in the absence of experimentally manipulated stimuli, which limits its interpretation with respect to information processing. Taken together, this body of research motivates the need for a methodological framework that classifies task-induced broadband EEG into periods of inhibition and excitation. This would greatly increase our understanding of the sequence of processing events that precede or follow a stimulus, allowing this activity to be related to other types of brain measurements, such as single/multiple units or neuroimaging recordings. In the current study, we expand on this idea in a paradigm that includes different phases of information processing performed by younger and older adults.

A fundamental step in classifying aperiodic activity into periods of inhibition and excitation is to establish *whether*, *when*, and *how* the appearance of a stimulus affects the ongoing aperiodic activity. However, a serious challenge to this endeavor is the need to separate the task-induced (non-phase-locked) aperiodic component from other task-evoked (phase-locked) EEG activity (i.e., event-related potentials, ERPs, in the time domain), both of which display a broadband distribution in the frequency domain. Gyurkovics et al. (2022) were the first to address this methodological issue using scalp EEG data collected from young adults. Their study showed reliable and systematic stimulus-induced changes in the aperiodic component, which were independent of the concurrent ERPs and scaled with the attentional demands of the task. The reported *stimulus-induced clockwise spectral rotations* are consistent with a decreased E:I ratio (increased inhibition) following stimulus onset and likely reflect a disruption of ongoing excitatory activity proportional to processing demands (Gratton, 2018; see also Zhang et al., 2023). However, the Gyurkovics et al. (2022) study was conducted using simple paradigms with minimal quantification of the participants’ performance, thus making it difficult to determine the behavioral consequences, if any, of the stimulus-induced spectral exponent shifts. Moreover, event-related spectrograms were quantified using a time window extending more than 1,000 ms, which precludes the detection of rapid changes in aperiodic activity accompanying information processing over time. Crucially, that study did not investigate the effects of age, which is expected to modulate the E:I balance. These three issues are addressed in the current study.

To summarize, the current study sought to determine the role of the aperiodic component – indexed by the spectral exponent – in the relationship between aging and cognitive processing while considering the temporal dynamics of this component. To this end, we analyzed scalp EEG data from younger and older adults performing a cued flanker task (**Fig. 1A**). We capitalized on changes in the aperiodic background activity induced by cues, which do not require any overt responses but provide information to prepare for the upcoming target stimuli (Bowie et al., 2021; Gratton et al., 1992). To capture the temporal dynamics of aperiodic activity, the cue-related EEG was divided into a pre-cue time window and three consecutive post-cue time windows (**Fig. 1B**). The pre-cue window, being free of any cue processing, served as a baseline. The three subsequent post-cue windows were intended to capture different phases of information processing (early, middle, and late).

**Figure 1.**
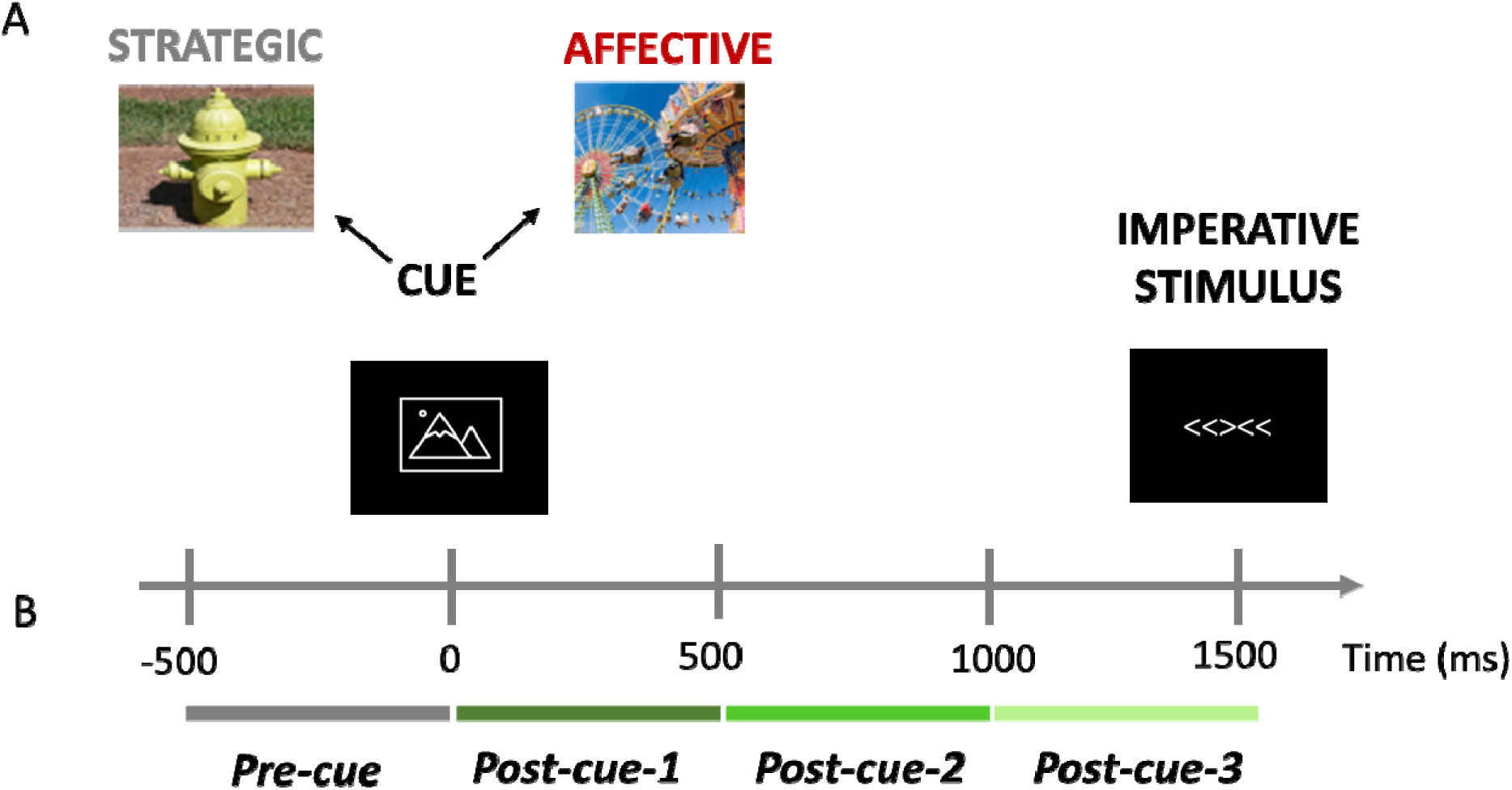
Behavioral task design and time windows for the EEG analyses. (A) Participants performed a cued flanker task. The warning cue presented at the beginning of the trial was followed by the imperative stimulus. Cues were images from the International Affective Picture System (Lang et al., 2008) and from an additional database (Iordan & Dolcos, 2015); images i this figure are for reference only. The cues were repetitive, task-relevant, and neutral (strategic blocks) or novel, task-irrelevant, and of variable valence (affective blocks); for details, see the text. (B) The cue-locked EEG recorded during the task was divide into four consecutive 500-ms time windows.

The results reveal hitherto unreported features of the aperiodic EEG, which allowed us to estimate shifts in the E:I balance as a function of the processing phase and age. As such, these findings expand our understanding of how dynamic neural communication shapes cognition in younger and older adults and have direct implications for neuroscientific models of cognitiv processing and age-related cognitive decline. Given that aperiodic neural activity is considered a key biomarker of healthy neural networks (Ahmad et al., 2022; Chini et al., 2022; Gao et al., 2017), this study could also have important implications for all neurocognitive domains examining normative and abnormal brain dynamics.

## 2. Methods

### 2.1. Participants

The study was conducted at the Beckman Institute of the University of Illinois at Urbana-Champaign. Forty-eight participants took part in the study. Two participants were excluded because spectral parameterization failed to provide converging solutions for them. Data from three additional participants had poor quality of the EEG power spectrum (the standard deviation, *SD*, of power across frequencies for each of these three participants was three times higher than the average *SD* of power across all participants). Although the inclusion/exclusion of these three participants did not significantly change the results, 1/*f^x^* analyses require data of the highest quality to be reliable, given that *all* frequencies (including those with very small power) are considered. Therefore, we limited the analyses presented in this article to the 43 participants with the highest data quality: 21 younger adults (mean age ± *SD* = 21.52 ± 2.82, 13 females) and 22 older adults (mean age ± *SD* = 71.23 ± 4.25, 10 females). The study was approved by the Institutional Review Board of the University of Illinois at Urbana-Champaign and followed the Declaration of Helsinki. Written informed consents were obtained from all participants. ERP analyses from a subset of these data, unrelated to the current report, were published by Bowie et al. (2021).

### 2.2. Experimental Task and Procedure

Participants performed a cued flanker task. The task design is presented in **Figure 1A**. The imperative stimulus array consisted of five horizontal arrows that were either congruent (e.g., <<<<<) or incongruent (e.g., <<><<). Participants indicated, as quickly and accurately as possible, which direction (left or right) the central (target) stimulus was pointing by pressing one of two keypads located on either side of the participant. Stimulus-response mapping was fixed (i.e., a left-pointing target stimulus always required a left-button press, and vice versa).

Images from the International Affective Picture System database (Lang et al., 2008) supplemented with images of neutral scenes from an additional database (Iordan & Dolcos, 2015) served as cues, preceding the presentation of the imperative stimulus array (their catalog numbers along with valence and arousal data are provided in the project repository at https://osf.io/dfbwa/). The pictorial cues were split into two sets to establish two different contexts for performing the flanker task: strategic and affective. In strategic blocks, cues were three neutral, low-arousal images (screw, fire hydrant, dresser), each of which indicated the probability of presenting a congruent stimulus array: Predict-Congruent had a *p*(congruent) of 75%; Predict-Incongruent had a *p*(congruent) of 25%; and No-Prediction had a *p*(congruent) of 50%. The cue types were equiprobable within each strategic block, and participants were explicitly told the probability of a congruent stimulus represented by each cue before commencing the task. Predict-Congruent and Predict-Incongruent images were counterbalanced across participants. In affective blocks, 288 images of varying arousal and valence served as cues. None of them indicated the probability of the imperative stimulus’ congruency. Instead, there were three task-irrelevant cue conditions that differed in valence while being equated in terms of arousal (low/high): Positive, Negative, and Neutral. All valence-arousal combinations were equiprobable and intermixed within a single affective block.

Each trial began with a 499-ms cue, followed by a 999-ms fixation period. Afterward, the imperative stimulus array appeared for 149 ms, followed by 1848 ms of fixation before the onset of the next trial. The response window began with the onset of the imperative stimulus and continued until the onset of the next trial. The imperative stimulus arrays were presented in white typeface on a black computer screen and subtended 2.23^◦^ × 0.46^◦^. Each cue overlaid a gray background with uniform dimensions such that each composite image subtended 6.98^◦^ × 5.35^◦^. All stimuli were presented on a monitor (19-inch CRT, refresh rate 60 Hz, screen resolution 1280 × 960; Dell Computer, Round Rock, TX, USA) using the E-Prime 2.0 software (Psychology Software Tools, Pittsburgh, PA, USA). Participants were seated 100 cm in front of a computer monitor centered at eye level.

There were three strategic blocks (288 trials each) and three affective blocks (288 trials each), yielding 1728 trials in total. The probability of a congruent trial within a single block was 50%. The strategic and affective blocks were alternated, and their order was counterbalanced across participants. All participants completed a set of practice trials prior to the task.

### 2.3. EEG Data Acquisition and Preprocessing

Scalp EEG was recorded from 59 Ag/AgCl active electrodes using a BrainAmp recording system (BrainVision Products). The electrodes were secured in an elastic cap according to the extended 10-20 international electrode placement system (Acharya et al., 2016). Horizontal and vertical electrooculograms (EOGs) were also recorded to monitor ocular artifacts. During recording, the data were filtered with a 0.10-250 Hz bandpass, digitized at a sampling rate of 500 Hz and referenced to the left mastoid. Impedance was kept < 10 kΩ.

The data were preprocessed using custom MATLAB 2022b codes (The MathWorks) incorporating EEGLAB 13.6.5 (Delorme & Makeig, 2004) and ERPlab 6.1.3 (Lopez-Calderon & Luck, 2014). The EEG was first re-referenced to the average mastoids and bandpass filtered with 0.5 and 50 Hz cut-off frequencies (to eliminate contamination from the power supply at 60 Hz). The data were then segmented into 2000-ms long epochs relative to the cue onset (−500 to 1500 ms). After excluding epochs with amplifier saturation and performing ocular correction (Gratton et al., 1983), epochs with peak-to-peak voltage fluctuations at any EEG channel exceeding 200 µV (600-ms window width, 100-ms window step) were discarded. Data from electrodes Fp1 and Fp2 were excluded as they often contain small residual ocular artifacts even after ocular correction. Epochs for which response latency in the preceding trial exceeded 1400 ms were also excluded, as late response-related activity from the previous trial could overlap with the baseline of the current trial, thus distorting the measurement of pre-cue activity. Since the accuracy of responses is not directly related to cue processing, epochs with both correct and incorrect responses were included^1^. The average number of artifact-free epochs per cue type across all participants was 221 (*SD* = 50, *min* = 73, *max* = 282).

### 2.4. Statistical Analyses

The data were analyzed and visualized in R 4.0.3 (R Core Team, 2021). *p-*values for *F-* tests were based on permutations for mixed ANOVA (Frossard & Renaud, 2021; Kherad-Pajouh & Renaud, 2015). We used 10,000 permutations, and the sign for a given parameter was reversed for a random half of the data points in each iteration (an equivalent approach was adopted in our previous work, Gyurkovics et al., 2022). Only planned comparisons were tested. *p-*values < 0.05 were considered significant. The materials, data, and R code for this project will be openly available in the project repository (https://osf.io/dfbwa/).

#### 2.4.1. Behavioral Analysis

Dependent variables (DVs) were mean reaction time (RT), mean error rate (ERR), and mean inverse efficiency score (IES, i.e., RT/*p*(correct); Townsend & Ashby, 1978). Fast guesses (i.e., RT ≤ 200 ms) and timeouts were discarded. Trials with incorrect responses were excluded from computing RT and IES. Since the EEG data were trimmed to epochs with RT < 1400 ms (for rationale, see section 2.3), this criterion was also applied to the behavioral data to maintain consistency across analyses. On average, 12% of trials (*SD* = 10%) were excluded, leaving approximately 1,520 trials per participant for analysis. The analyses replicated previously reported effects (Bowie et al., 2021; Gratton et al., 1992), indicating that data trimming did not impact the results.

To evaluate results within the strategic and affective contexts separately (within-context ANOVAs, hereafter), DVs were subjected to mixed ANOVAs with Age Group as a between-subject factor (younger, older) and two within-subject factors: Congruency (congruent, incongruent) and Cue Type (predict-congruent, no-prediction, predict-incongruent, for the strategic context; positive, neutral, negative, for the affective context). To compare results across task contexts, data were collapsed across task contexts, and the cue type factor was replaced with the within-subject Task Context factor (strategic, affective) (between-context ANOVA, hereafter).

#### 2.4.2. Spectral Analysis

To investigate the temporal variation of the aperiodic component, the cue-locked epochs were divided into four successive time windows of equal length, representing, respectively, pre-cue activity (−501 – −1 ms; pre-cue/baseline), activity directly after the cue (0 – 500 ms; post-cue-1), mid-interval activity (500 – 1000 ms; post-cue-2), and activity directly before the target stimulus (1000 – 1500 ms; post-cue-3) (see **Fig. 1B**). Single-trial total power spectra were then computed for each time window, electrode, and participant, using MATLAB’s built-in fast Fourier transform (FFT) function. Before FFT, the signal was zero-padded to 256 points to ensure that signal length was a power of 2 for the FFT. The spectral resolution was 1.95 Hz. Frequencies < 1.95 and > 44.92 Hz were removed to avoid frequencies whose power estimates were based on < 2 cycles and to ensure frequencies affected by the low-pass filter were omitted. The resulting total power spectra were then averaged across trials for each time window (pre-cue, post-cue-1, post-cue-2, post-cue-3), EEG channel (57 in total after excluding Fp1 and Fp2), and cue type (Predict-Congruent, No-Prediction, Predict-Incongruent, Positive, Neutral, Negative) within each participant separately. To account for the presence of ERPs in the post-cue windows, the spectra of the ERPs (i.e., the cross-trial time-domain averages) were also quantified for each time window × electrode × cue type × participant. These spectra were then subtracted from the total power spectra to yield power spectra after ERP removal, using the procedure described by Gyurkovics et al. (2022).

To separate oscillatory and aperiodic spectral components, single-electrode power spectra before and after ERP removal were then parametrized using the *specparam* algorithm (version 1.0.0; Donoghue et al., 2020) with the following settings: peak width limits = 3-8 Hz; the maximum number of peaks = 3; peak threshold = 2 *SD*; aperiodic mode = ‘fixed’. These parameters were determined on the basis of a preliminary analysis on a random sample of 20 participants, following guidelines by Ostlund et al. (2022). The aperiodic component at each electrode for each participant and time window was then reconstructed in linear space as 10^(*ß*+*x*log10(*f*))^, where *ß* is the offset in log space, *f* is frequency, and *x* (with a negative sign) is the exponent. The exponent values were retained for further analyses, with more negative values indexing steeper spectra (clockwise rotation) and decreased E:I ratio (increased inhibition).

The quality of spectral parametrization was assessed using *specparam*’s model *R^2^*. Since 14 parieto-temporal electrodes near the edge of the electrode cap showed relatively poorer fit (median of participants’ average *R^2^* < 0.90 for any time window × cue type), they were excluded from all analyses. Their reduced fit was likely due to muscle artifacts, affecting the estimation of high-frequency power. To balance the statistical power of the different levels of the electrode cluster factor, the four outermost parietal electrodes (P7, P8, PO7, PO8) were also excluded. The remaining 39 electrodes with satisfactory fit are shown in **Figure 4B**. Average *R^2^*s were 0.95 (*SD* = 0.05) for the younger group and 0.93 (*SD* = 0.03) for the older group. While younger participants showed a relatively higher *specparam R^2^* than older adults [*F*(1,41) = 4.05, *p* = 0.05], the fit was satisfactory in both age groups.

Given the novelty of the procedures used by Gyurkovics et al. (2022) to remove the ERP spectra, we first performed two auxiliary analyses to replicate their findings. First, to examine whether the ERPs contributed to the cue-locked background activity, the exponents estimated on the spectra after ERP removal were compared with those estimated on the spectra before ERP removal. Second, to assess whether the cue induced a change in the aperiodic component (*cue-induced spectral shift*, hereafter), post-cue exponents after ERP removal were compared against the pre-cue exponents. These analyses were performed on the exponent values averaged across electrodes and cue types for each time window separately using a series of one-way within-subject ANOVAs.

As the pre-cue window served as a baseline in the analyses, we also tested whether the pre-cue exponent (averaged across electrodes) showed any within-subject effects of Cue Type or Task Context that could obscure the experimental effects in the post-cue period. The between-subject Age Group factor was also included to assess age-related changes in baseline aperiodic activity.

Afterward, we analyzed the temporal dynamics and effects of experimental manipulation on cue-induced spectral shifts. To this end, the post-cue spectral exponents after ERP removal in each of the three post-cue windows (i.e., post-cue-1, post-cue-2, and post-cue-3) were subtracted from the pre-cue exponent for each electrode × cue type × participant, yielding Shift1, Shift2, and Shift3, respectively. These cue-induced spectral shifts were then subjected to the within-context and between-context ANOVAs, all of which included Age Group as a between-subject factor and two within-subject factors: Cue Type/Task Context and Time Window (Shift1, Shift3). Cue Type and Task Context levels were the same as in the behavioral analyses. Shift2 was deliberately excluded from these analyses as we did not observe a significant group-level Shift2 (see section 3.2). To investigate possible differences in scalp distribution, the data were averaged over two electrode clusters covering fronto-central and centro-parietal regions (**Fig. 4B**), constituting an additional within-subject factor in these analyses.

#### 2.4.3. Neuro-Behavioral Correlations

Multiple rank-based regression – a non-parametric, robust alternative to the traditional likelihood or least-squares estimators (Kloke & Mckean, 2012) – was used to determine the effects of aging and aperiodic activity on overall performance (indexed by the IES) and magnitude of the congruency effect (indexed by incongruent minus congruent IES), for each time window separately (pre-cue, post-cue-1, post-cue-2, post-cue-3). The *simple model* included the one of the aperiodic predictors (pre-cue exponent, Shift1, Shift2, or Shift3, depending on the time window), whereas the *additive model* additionally included continuous age. Although Shift2 was excluded from the ANOVAs, as there was no significant difference between post-cue-2 and pre-cue exponents (see section 3.2), we chose to re-include it in the correlation analyses. This is because a non-significant group-level effect might reflect large inter-individual variability in the post-cue-2 window, which could be potentially interesting for an individual-difference perspective.

The model including the interaction between the predictors was discarded as it did not perform better than the additive model for any DV in any time window (non-significant dispersion-reduction tests, an equivalent of χ*^2^* in classic regression; *F*s ≤ 3.81). Since the effects showed relatively widespread scalp distributions and analyses for strategic and affective contexts produced largely consistent results, the statistics are reported for the data averaged across all 39 electrodes and both task contexts. For visualization purposes, the figures present the regression *beta* estimates on single electrodes. All variables were mean centered prior to these analyses.

## 3. Results

### 3.1. Contextual Variability Supports the Behavioral Performance of Older Adults

RT, ERR, and IES were subjected to mixed ANOVAs to test the experimental effects of Age Group, Congruency, and Cue Type/Task Context. Since the results were largely consistent across all DVs, the statistics are reported for IES only, as this DV combines both speed and accuracy information, hence providing a robust summary of performance (Townsend & Ashby, 1978). **Figure 2A** presents an overview of the behavioral results.

**Figure 2.**
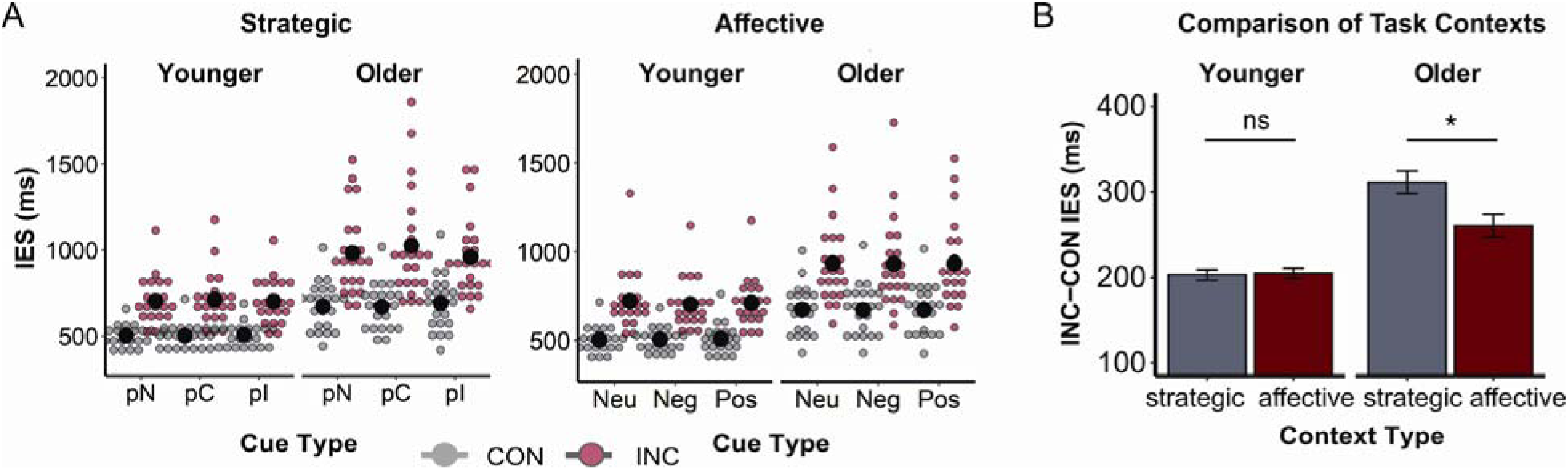
Behavioral results. (A) Inverse efficiency scores (IES) in milliseconds (ms) for the strategic context (left) and affective context (right). Black circles depict means across participants by cue type and congruency. Colored dots represent individual participants’ scores for the congruent (gray, CON) and incongruent (red, INC) conditions. *pN,* No-Prediction; *pC*, Predict-Congruent; *pI*, Predict-Incongruent; *Neu,* Neutral; *Neg,* Negative; *Pos*, Positive. (B) Congruency effect (INC-CON) in mean inverse efficiency score (IES) in milliseconds (ms) by task context and age group. Bars depict the mean across participants ± within-subject standard error; *ns*, non-significant; *\**, *p* < 0.05.

The analysis for the strategic context replicated previously reported effects (Bowie et al., 2021). Participants were more efficient in the congruent vs. incongruent condition [*F*(1,41) = 84.04, *p* < 0.001, η*p²* = 0.67], and older adults were less efficient than younger adults [*F*(1,41) = 28.85, *p* < 0.001, η*p²* = 0.41]. Moreover, Congruency interacted with Cue Type in the strategic context [*F*(2,82) = 8.76, *p* < 0.001, η*p²* = 0.18]. Performance was lower in the congruent condition when the incongruent condition was predicted compared to when a congruent stimulus was expected [*t*(42) = 2.25, *p* = 0.03, *d* = 0.34], or no congruency prediction could be made [*t*(42) = 2.44, *p* = 0.02, *d* = 0.37]. Conversely, performance was lower in the incongruent condition when the congruent condition was predicted compared to when an incongruent stimulus was expected [*t*(42) = 2.72, *p* = 0.01, *d* = 0.41] or congruency could not be predicted [*t*(42) = 2.08, *p* = 0.04, *d* = 0.32]. While the effects of Congruency and Age Group were replicated in the affective context [*F*(1,41) = 89.70, *p* < 0.001, η*p²* = 0.69, and *F*(1,41) = 20.96, *p* < 0.001, η*p²* = 0.34, respectively], there were no effects of Cue Type (*F*s ≤ 1.43).

A between-context ANOVA was performed to disentangle the global impact of strategic cues (which were neutral images repeated over trials) and affective cues (which varied in valence and were unique on each trial within a block). The analysis replicated the Congruency and Age Group effects described above [*F*(1,41) = 89.32, *p* < 0.001, η*p²* = 0.69, and *F*(1,41) = 25.34, *p* < 0.001, η*p²* = 0.38, respectively]. We also observed a significant Task Context effect [*F*(1,41) = 4.97, *p* = 0.03, η*p²* = 0.11], which was qualified by Age Group [*F*(1,41) = 8.61, *p* < 0.001, η*p²* = 0.17] and Congruency [*F*(1,41) = 5.61, *p* = 0.02, η*p²* = 0.69]. Interestingly, there was also a three-way interaction between these factors [*F*(1,41) = 6.42, *p* = 0.01, η*p²* = 0.14]. While younger participants did not differ significantly in the congruency effect (incongruent *minus* congruent) between task contexts [*t*(20) = 0.20, *p* > 0.05], older participants demonstrated a reduced congruency effect in the affective vs. strategic context [*t*(21) = 2.74, *p* = 0.01, *d* = 0.58] (**Fig. 2B**), that was driven by their faster and more accurate responses in the affective-incongruent vs. strategic-incongruent condition [*t*(21) = 3.19, *p* < 0.001, *d* = 0.68]. Consequently, there was no significant between-group difference in the congruency effect in the affective context [*t*(28) = 1.15, *p* > 0.05].

To investigate why older adults performed better in the affective compared to the strategic context, we tested the Age Group × Task Context × Congruency interaction on trials with neutral cues only (‘no-prediction’ cues from the strategic context and neutral cues from the affective context). These cues differed in terms of novelty (same on every trial in a strategic block vs. unique on every trial in an affective block) but were comparable in terms of valence (all neutral) and task relevance (all unpredictive). A three-way interaction was observed for this limited (neutral only) cue set [*F*(1,41) = 6.72, *p* = 0.01, η*p²* = 0.14], bolstering the interpretation that the greater contextual variability and novelty introduced by repeatedly changing cues in the affective context supports the cognitive performance of older adults. This interpretation is further corroborated by the absence of significant effects of cue valence in the affective ANOVA (see above), as well as the absence of block order or arousal effects in the follow-up analyses (see also Footnote 2 in Bowie et al., 2021).

### 3.2. Cue-Related Changes in Aperiodic Background Activity above and beyond the Contribution of ERPs

Consistent with our previous work (Gyurkovics et al., 2022), the exponent values were significantly reduced (i.e., were less negative, flatter spectrum) when the frequency spectrum of the ERPs was removed in each of the three post-cue windows [*F*(1,42) ≥ 76.18, *p* < 0.001, η*p²* ≥ 0.64], indicating that the ERPs contribute to the shape of the event-locked EEG spectrum and must be removed before estimating aperiodic parameters (**Fig. 3A**). Further analyses focused on the post-cue estimates after ERP removal (**Fig. 3B-C**).

**Figure 3.**
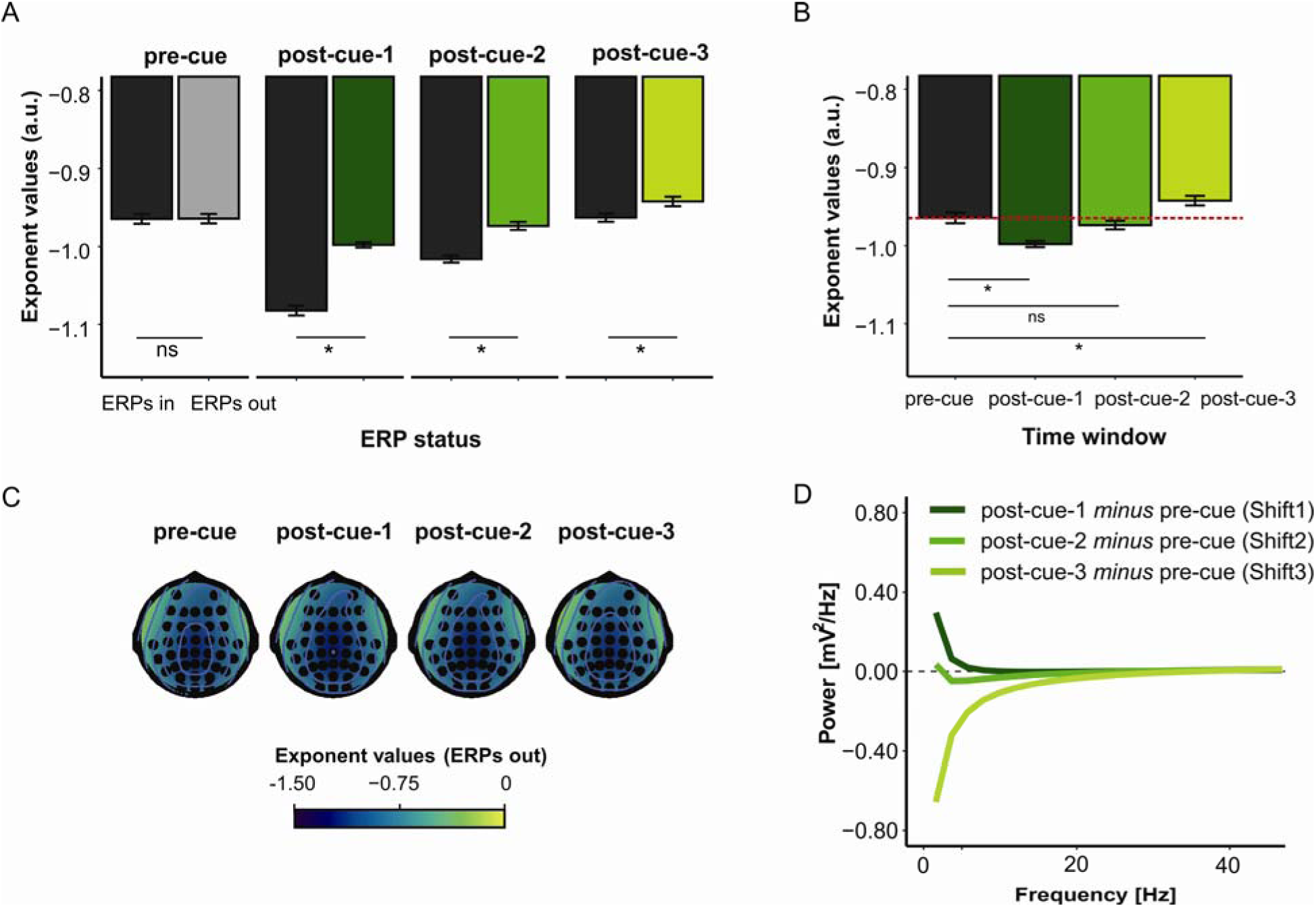
Aperiodic component overview. (A) Average exponent before (black) and after (gray/green) removal of the ERP spectrum (ERPs in and ERPs out, respectively) by time window. (B) Average exponent by time window. The red dashed line indicates the mean value of the pre-cue/baseline period. (C) Scalp distribution of the absolute exponent values in each time window. (D) Cue-induced spectral shifts (post-cue aperiodic components after subtracting pre-cue/baseline component) across frequencies in each time window, termed *Shift1*, *Shift2*, and *Shift3*. For both (A) and (B), error bars depict the mean across participants ± within-subject standard error. *ns*, nonsignificant, *\**, *p* < 0.05. For (B), (C), and (D), the post-cue exponent values are after ERP removal. For all panels, more negative values indicate steeper spectra.

Cue-induced exponent changes were observed in two of the three post-cue windows. Compared to the pre-cue window, the post-cue-1 exponent was more negative, indicating a clockwise rotational shift [Shift1; *F*(1,42) = 28.48, p < 0.001, η*p²* = 0.40], whereas the post-cue3 exponent was less negative, indicating a counterclockwise shift [Shift3; *F*(1,42) = 4.61, p = 0.04, η*p²* = 0.10]. The absence of a significant difference between the pre-cue and post-cue-2 exponents indicates that there was no detectable group-level shift in the mid-interval, relative to the pre-cue period [Shift2; *F*(1,42) = 0.89, *p* > 0.05].

Considering the pre-cue (baseline) activity, no effects of Cue Type or Task Context were found [*F*s < 1], indicating that the pre-cue activity provided an unbiased baseline for post-cue comparisons. At the same time, consistent with research showing an age-related decrease in ongoing (baseline) aperiodic activity (for a review, see Ostlund et al., 2022), the pre-cue exponent was less negative for older compared to younger adults [*F*(1,41) ≥ 21.43, *p <* 0.001, η*p²* ≥ 0.34] indicating a flatter power spectrum in the former age group.

### 3.3. Dynamic Nature of Aperiodic Background Activity and Age-Related Changes

**Figures 3D** and **4A** present an overview of cue-induced spectral shifts, referred to as Shift1 (post-cue-1 *minus* pre-cue), Shift2 (post-cue-2 *minus* pre-cue), and Shift3 (post-cue-3 *minus* pre-cue). These spectral shifts were subjected to strategic, affective, and between-context ANOVAs. As mentioned, Shift2 was excluded, as we did not find a group-level exponent difference in the post-cue-2 vs. pre-cue comparison. Statistics are shown in **Table 1**. Since Age Group and Time Window effects were consistent across these analyses, the follow-up tests are reported for the between-context ANOVA only.

**Figure 4.**
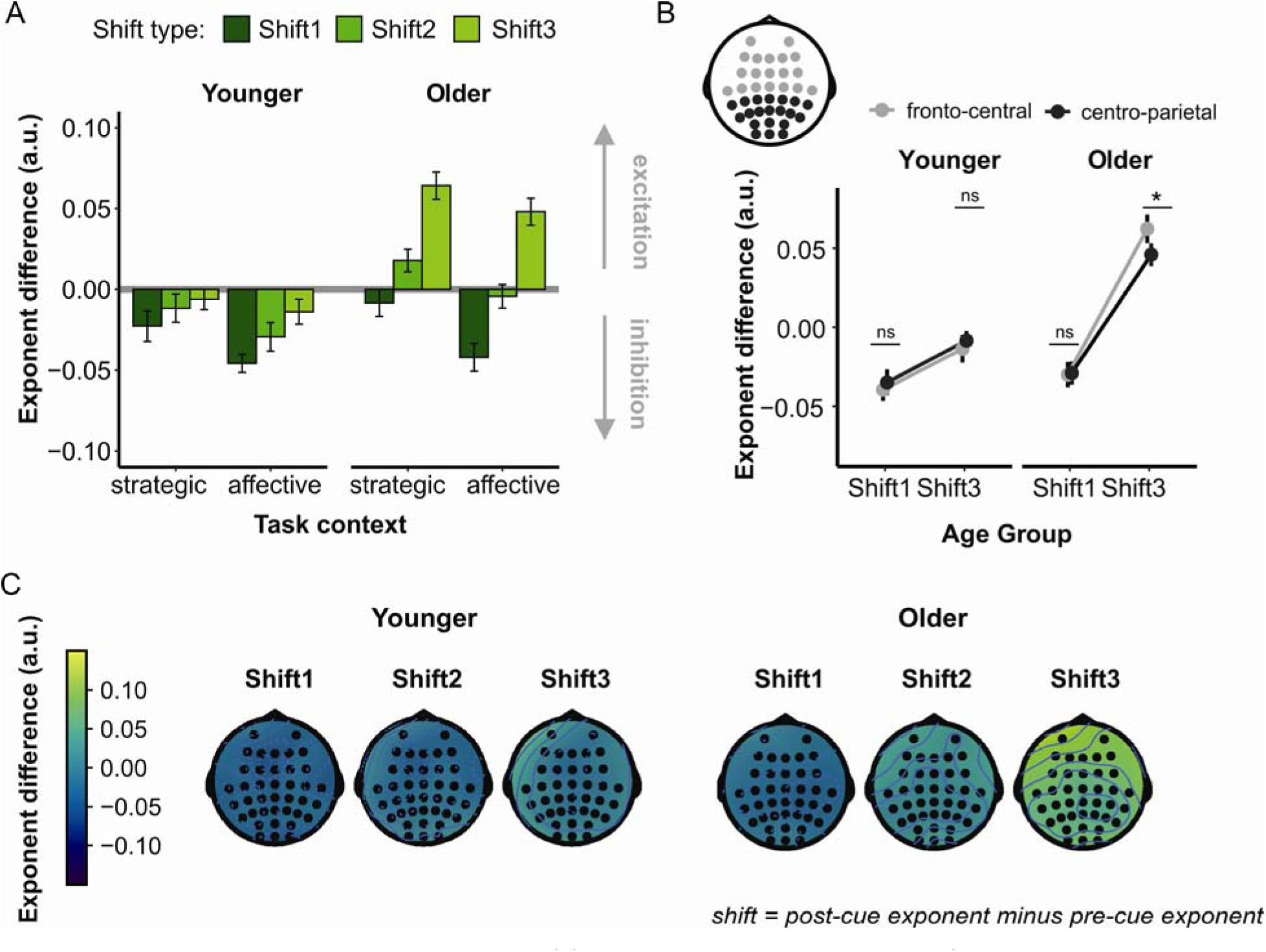
Dynamic nature of the aperiodic component. (A) Average cue-induced spectral shifts (post-cue after the ERP removal *minus* pre-cue exponent) by age group, task block, and time window, termed *Shift1*, *Shift2*, and *Shift3*. Error bars depict the mean across participants ± within-subject standard error. (B) Average cue-induced spectral shifts by age group, time window, and electrode cluster. Fronto-central (light gray) and centro-parietal (dark gray) electrode clusters are depicted on the scalp above the line plot. *ns*, non-significant, *\**, *p* < 0.05. (C) Scalp distribution of the cue-induced spectral shifts by time window for younger (left) and older participants (right). For all panels, more negative values indicate steeper spectra.

**Table 1.** Summary of ANOVA Results for Between-Context, Strategic, and Affective Effects

| Effects | *Between-Context* |  |  | *Strategic* |  |  | *Affective* |  |  |
| --- | --- | --- | --- | --- | --- | --- | --- | --- | --- |
| | *F* | *p* | $\eta p^2$ | *F* | *p* | $\eta p^2$ | *F* | *p* | $\eta p^2$ |
| Age Group | 7.85 | 0.01 | 0.16 | 12.34 | 0.00 | 0.23 | 2.81 | 0.09 | --- |
| Task | 13.58 | 0.00 | 0.25 | 0.71 | 0.49 | --- | 2.08 | 0.12 | --- |
| Age Group × Task | 0.77 | 0.40 | --- | 0.80 | 0.46 | --- | 0.95 | 0.38 | --- |
| Time | 59.13 | 0.00 | 0.59 | 34.34 | 0.00 | 0.46 | 50.29 | 0.00 | 0.55 |
| Age Group × Time | 17.31 | 0.00 | 0.30 | 13.59 | 0.00 | 0.25 | 12.63 | 0.00 | 0.24 |
| Cluster | 0.00 | 0.95 | --- | 1.00 | 0.32 | --- | 0.87 | 0.33 | --- |
| Age Group × Cluster | 2.72 | 0.11 | --- | 5.09 | 0.03 | 0.11 | 0.66 | 0.41 | --- |
| Task × Time | 4.49 | 0.04 | 0.10 | 0.19 | 0.82 | --- | 2.42 | 0.08 | --- |
| Age Group × Task × Time | 0.02 | 0.90 | --- | 1.22 | 0.30 | --- | 1.34 | 0.25 | --- |
| Task × Cluster | 3.31 | 0.08 | --- | 2.51 | 0.09 | --- | 0.42 | 0.66 | --- |
| Age Group × Task × Cluster | 0.91 | 0.36 | --- | 1.32 | 0.27 | --- | 0.00 | 1.00 | --- |
| Time × Cluster | 6.20 | 0.02 | 0.13 | 6.21 | 0.02 | 0.13 | 2.18 | 0.13 | --- |
| Age Group × Time × Cluster | 5.98 | 0.02 | 0.13 | 3.28 | 0.08 | --- | 5.07 | 0.02 | 0.11 |
| Task × Time × Cluster | 1.06 | 0.31 | --- | 1.07 | 0.35 | --- | 0.24 | 0.79 | --- |
| Age Group × Task × Time × Cluster | 0.04 | 0.84 | --- | 0.21 | 0.82 | --- | 0.14 | 0.87 | --- |

*Note.* Task refers to the task context (levels: strategic, affective) or to the strategic/affective cue type (levels: predict-congruent, no-prediction, predict-incongruent/positive, neutral, negative); degrees of freedom (*dfs*) for all effects are (1,41) except for task effects in the strategic context and affective context, for which *dfs* are (2,82).

All analyses showed a significant Age Group effect. Compared to younger adults, older adults demonstrated a less negative spectral shift, indicating a counterclockwise spectral rotation and an increased E:I ratio. A Time Window effect was also significant across all analyses, indicating changes in the time course of cue-locked aperiodic activity. The initially negative spectral shift (signifying a clockwise spectral rotation and a decreased E:I ratio for Shift1) decreased over time to become a positive spectral shift before the target appeared (counterclockwise rotation and an increased E:I ratio for Shift3). Moreover, Age Group interacted significantly with Time Window across all analyses (**Fig. 4A**). Interestingly, Shift1 did not differ between younger and older participants (no significant age-group difference in E:I ratio), *t*(38.67) = 0.61, *p* > 0.05. Instead, what differentiated older from younger adults was their greater counterclockwise rotation in time (increased E:I ratio and decreased inhibition), which emerged as a significant Age Group difference for Shift3, *t*(39.77) = 3.56, *p* < 0.001, *d* = 1.08.

The interaction between Age Group and Time Window was further qualified by significant effects for Electrode Cluster in the between-context and affective-context analyses. While most spectral shifts showed a widespread distribution in both age groups (no significant differences between fronto-central and centro-parietal clusters, *t* ≤ 0.69, *p* > 0.05), Shift3 was larger at the fronto-central than centro-parietal cluster in older adults, *t*(21) ≥ 2.55, *p* < 0.01, *d* ≥ 0.54 (**Fig. 4B**). Collectively, these results point to the dynamic nature of stimulus-related changes in the aperiodic component, indicating that the feature distinguishing older from younger adults is a greater counterclockwise power redistribution over time. This indicates an increasing E:I ratio and decreasing inhibition with advancing age.

### 3.4. Attention-Dependent Changes in the Aperiodic Background Activity

Although the ANOVAs did not show any Cue Type effects when strategic and affective contexts were considered separately (see **Table 1**), we did observe a significant Task Context effect in the between-context ANOVA, with a more negative spectral shift (i.e., a more clockwise spectral rotation indicating increased inhibition) in the affective compared to the strategic context. This effect was not qualified by Age Group in the between-context comparison. However, a significant Age Group × Task Context interaction was observed when comparing the *neutral* cues from the two task contexts [*F*(1,41) = 6.71, *p* < 0.001, η*p²* = 0.50]. While the spectral shift was less negative in response to repetitive neutral cues in older compared to younger adults (strategic context: *t*(35.48) = 3.20, *p* < 0.001, *d* = 0.98), there was no significant difference between the age groups in response to more novel neutral cues (affective context: *t*(35.73) = 0.64, *p* > 0.05; see **Fig. 4A**). This suggests that the difference in the cue-induced spectral shift between younger and older adults diminished in response to the more novel cues presented in the affective context. This effect is consistent with the behavioral data, showing improved performance in the affective context in older adults, and further indicates that increased contextual variability may support cognitive performance in older adults.

### 3.5. Neuro-Behavioral Relationships

To further understand the relationships between aging, aperiodic activity, and behavior, we performed a series of neuro-behavioral correlations (**Figs. 5** and **6**). Consistent with the ANOVA results, older age was associated with a less negative pre-cue (baseline) exponent, further supporting the notion that aging is accompanied by an overall increase in the E:I ratio, indicating reduced inhibitory function (Ostlund et al., 2022; Voytek & Knight, 2015). Interestingly, however, when considering the cue-induced spectral shifts, age significantly correlated with Shift2 and Shift3 but not Shift1 (**Fig. 5A**). These results complement the ANOVA findings (significant age-group effect for Shift3 but not Shift1), further suggesting that the initial aperiodic response to the cue (indexed by Shift1) was not associated with age-related changes in the E:I balance, and that age only began to contribute to the cue-induced spectral shift after some time (middle and late post-cue periods in this study, indexed by Shift2 and Shift3, respectively).

**Figure 5.**
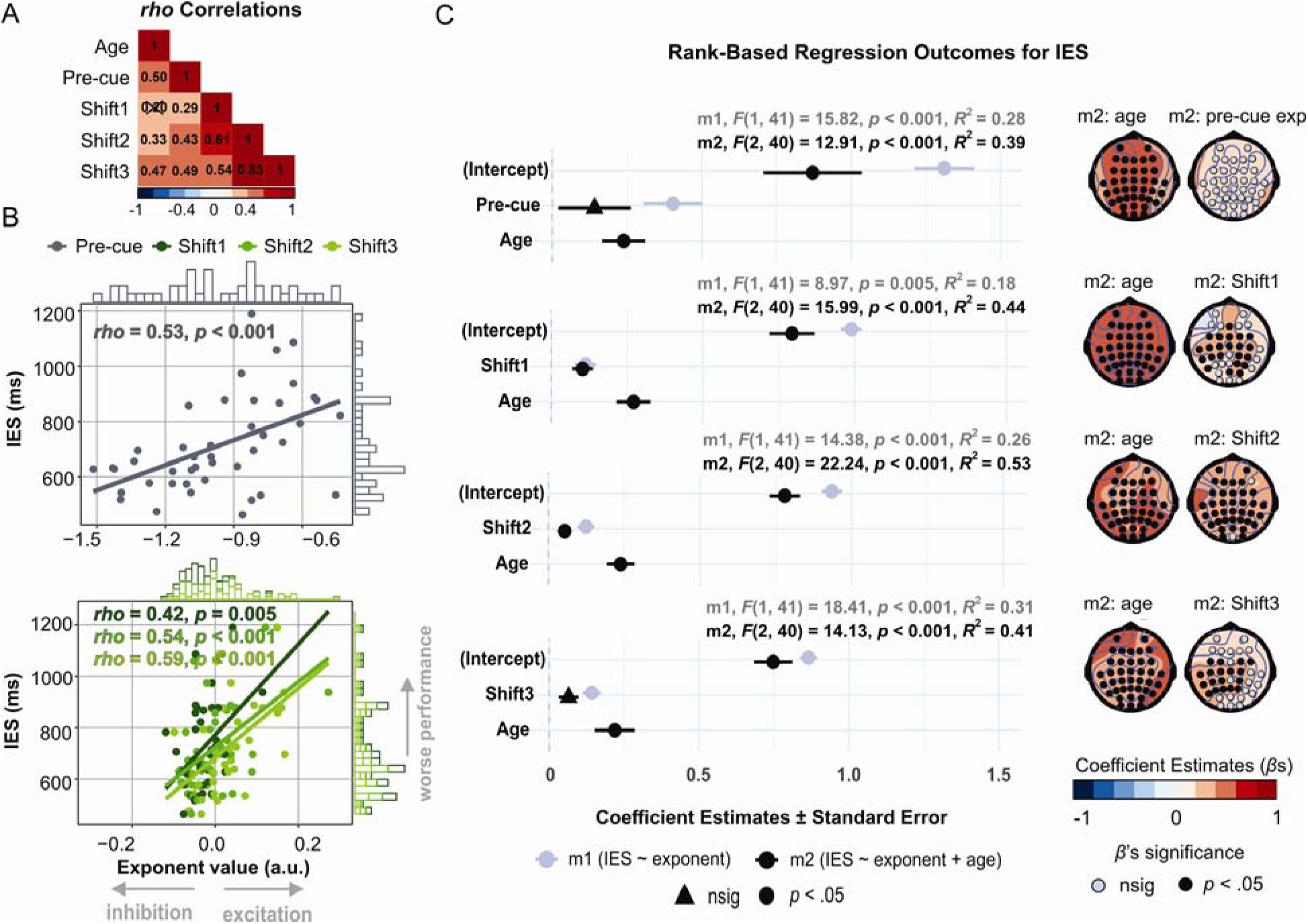
Overall performance as a function of the spectral exponent. (A) Spearman’s *rho* correlations between continuous age, pre-cue exponent, and post-cue spectral shifts (post-cue exponent after subtracting the pre-cue value). Non-significant estimates (*p* > 0.05) were crossed out. (B) Inverse efficiency score (IES) in milliseconds (ms) as a function of absolute pre-cue exponent (upper) and spectral shifts (post-cue exponents after subtracting the pre-cue exponent; lower). Coefficients are Spearman’s *rho*s. (C) Outcomes of rank-based regressions for each time window. The left panel shows regression coefficients (*beta*s, βs) ± standard error for the simple model (m1, gray) and additive model (m2, black). The right panel displays the scalp distribution of regression coefficients (βs) for the effects of age and exponent from m2 (*p*-values on the scalp maps uncorrected for multiple comparisons).

**Figure 6.**
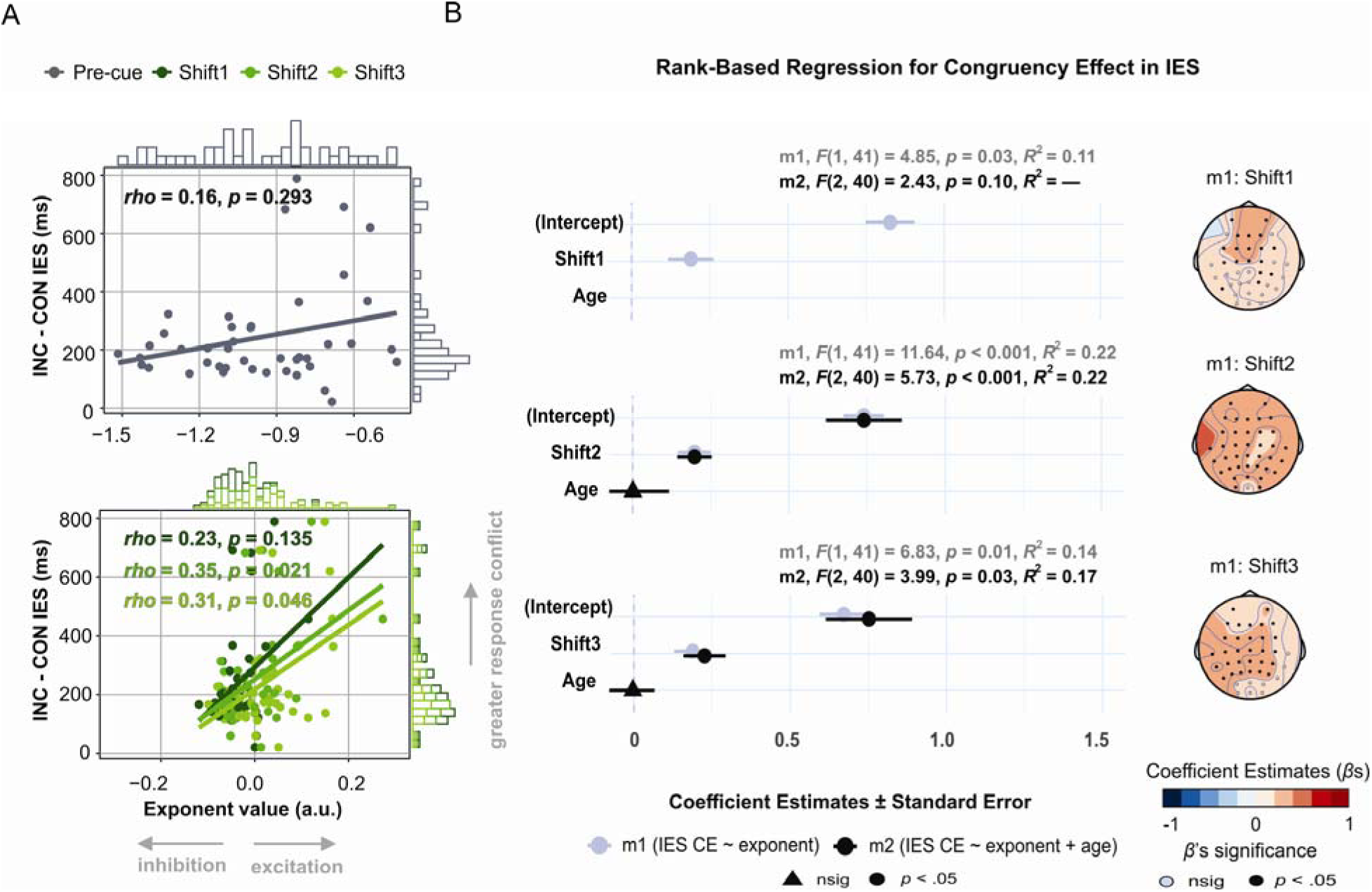
The magnitude of the congruency effect as a function of the spectral exponent. (A) Congruency effect (INC-CON) in inverse efficiency score (IES) in milliseconds (ms) as a function of absolute pre-cue exponent (upper) and spectral shifts (post-cue exponents after subtracting the pre-cue exponent; lower). Coefficients are Spearman’s *rho*s. (B) Outcomes of rank-based regressions for each post-cue window (note that the models with the pre-cue exponent were non-significant and are therefore omitted). The left panel shows regression coefficients (*beta*s, βs) ± standard error for the simple model (m1, gray) and additive model (m2, black). The right panel displays the scalp distribution of regression coefficients (βs) for the effect of exponent from m1 (*p*-values on the scalp maps uncorrected for multiple comparisons).

To determine how the aperiodic activity and aging contributed to overall performance (indexed by IES) and to the congruency effect (incongruent *minus* congruent IES), we fit a series of rank-based regression models. Regarding overall performance (**Fig. 5B-C**), a simple model including the pre-cue exponent or post-cue shift as a predictor of IES was significant across all time windows, indicating that the more negative the baseline exponent or the more negative the post-cue shifts (all indicating increased inhibition), the higher the task performance. Models including age as an additional predictor (additive models, hereafter) were also significant in each time window and indicated that older age was associated with lower behavioral outcomes. Importantly, when age was added to the models, the pre-cue exponent and Shift3 no longer significantly predicted IES, indicating that their associations with IES observed in the simple models could be fully explained by age-related changes in aperiodic activity. At the same time, Shift1 and Shift2 continued to be significant predictors after the addition of age, indicating that the cue introduced changes in the aperiodic activity that predicted subsequent performance regardless of age.

Regarding the efficiency of resolving the flanker-induced response conflict, indexed by the congruency effect (**Fig. 6**), a simple model including the post-cue shift as a predictor of the congruency effect in IES was significant across all post-cue time windows: the more negative the post-cue shift (i.e., the greater the shift towards inhibition), the smaller the subsequent congruency effect. The additive model was significant for Shift2 and Shift3, but the age effect was non-significant. Neither model was significant for the pre-cue window [*F*s < 1], indicating that baseline aperiodic activity, similarly to age, is unrelated to the magnitude of the congruency effect.

In summary, the analyses reported here suggest that aperiodic neural activity substantially affects subsequent performance on the flanker task. While age contributed to overall performance, cue-induced spectral shifts were not only able to predict overall performance but also the magnitude of the congruency effect. Furthermore, results suggest that the cue-induced spectral shift is a mixture of age-dependent and age-independent processes, whose relative contribution to performance depend on the information processing timescale (**Fig. 5B**). Specifically, since Shift1 did not correlate with age and the addition of age to the IES model hardly changed its estimate (△β = 0.009), the spectral shift immediately following the cue appears to reflect age-invariant stimulus processing. Conversely, the relationship between the latest shift (Shift3) and overall performance was canceled out when age was added to the IES model, indicating that it was entirely driven by age-dependent changes in information processing. In line with this logic, age-dependent and age-invariant stimulus processing co-contributed to the shift in the mid-interval, as shown by the additional model in which Shift2 still significantly predicted IES after regressing out either age or Shift1 (the latter representing age-invariant stimulus processing). Yet, the explained variance substantially dropped in both cases (△*R^2^* = 17% and 15%, respectively). Notably, after regressing out both age and Shift1, this model ceased to be significant [*F*(1,41) = 1.84, *p* > 0.05], indicating that there was no additional variance in Shift2 that would explain IES over and above the effect of age-dependent and age-invariant cue processing present in the first time window.

## 4. Discussion

This study provides an in-depth analysis of stimulus- and age-related changes in the spectral exponent, an overarching measure of aperiodic background neural activity, indicating rotational shifts in broadband power. To this end, we analyzed scalp-recorded EEG data from younger and older adults who completed a cued flanker task. In this task, the pictorial cues were either repetitive, neutral, and task-related (creating a strategic context) or relatively novel, of varying valence, and unrelated to the task (affective context). This study extends our knowledge of stimulus-induced changes in the spectral exponent (see Gyurkovics et al., 2022) by showing that cues, signaling upcoming targets, trigger systematic changes in EEG background activity independently from the ERPs elicited by the same stimuli. In addition to the experimental effects, we also observed significant individual variations in the exponent in relation to age, stimulus processing phase, and subsequent behavioral performance. Collectively, the findings extend our current knowledge of the neural dynamics underlying aging and cognitive processing and bring these phenomena together within a unified framework.

### 4.1. Contextual Variability Supports the Cognitive Functioning of Older Adults by Altering Aperiodic Neural Activity

The behavioral analyses revealed some novel, hitherto unreported findings: overall, performance was higher when pictorial cues were relatively novel (affective context) than when they were repeated (strategic context). At the same time, cue valence itself had no detectable effect on behavior. This novelty (task-context) effect was further qualified by age group, indicating that presenting relatively more novel and varied cues made older adults more efficient and, thus, behaviorally more comparable to younger adults (**Fig. 2B**). The data we report, therefore, suggest that the presentation of relatively novel and variable pictorial cues created a task context that helped older adults to maintain increased engagement throughout the task, which consequently resulted in their more efficient performance. In contrast, younger adults were able to maintain high-level performance regardless of cue characteristics.

The EEG data showed systematic cue-induced changes in the ongoing background aperiodic activity (i.e., cue-induced spectral shifts) that varied depending on the task context. Specifically, the cue induced a more pronounced clockwise rotation (i.e., more negative post-cue vs. pre-cue exponent in log-log space) in the affective than in the strategic context. Consistent with previous findings (Gyurkovics et al., 2022), this task-context effect indicates increased inhibition in more novel and variable settings, which require more frequent updating of active representation status (Gratton, 2018; see also Zhang et al., 2023). Interestingly, the observed task-context effect was further qualified by age group when only neutral cues were considered. There was a significant age group difference in the spectral shift for repeated neutral cues used in the strategic context but not for the more novel and variable neutral cues used in the affective context. These findings are consistent with the behavioral results and suggest that the relatively greater inhibition induced by more novel cues helped older participants to overcome, at least in part, the age-related E:I imbalance towards excitation (Merkin et al., 2023; Ostlund et al., 2022; Thuwal et al., 2021; Voytek & Knight, 2015), making their cue-induced aperiodic response, as well as their subsequent performance, more comparable to that of younger adults. Consistent with our previous work (Gyurkovics et al., 2022), the observed cue-induced spectral shifts in the aperiodic component showed broad scalp distributions (no significant effects of electrode cluster were observed), further suggesting that the alternations in the E:I balance involve widespread changes in cortical activity.

### 4.2. Dynamics of Aperiodic Neural Activity and Their Consequences for Behavior

This study allowed us to examine the temporal dynamics of the aperiodic component related to different phases of stimulus processing. The results revealed that, compared to the pre-cue (baseline) period, the cue initially induced a clockwise shift in the ongoing power spectrum (Shift1), which became counterclockwise over time (Shift3), pointing to the transient nature of the aperiodic neural activity. Notably, there was no difference between age groups in the early phase of cue processing (Shift1). However, older adults (compared to younger adults) demonstrated a greater counterclockwise rotation in the late processing phase (Shift3) (**Fig. 4**). These experimental findings were further supported by significant correlations between age and spectral shifts in the middle and late but not early processing phase (**Fig. 5A**).

Similarly to the task-context effects discussed in the previous section, the observed temporal effects can also be explained within the E:I balance framework. The clockwise rotation immediately following the cue (Shift1) is consistent with a shift towards inhibition that temporarily halts ongoing processing to allow for new representations to be established (Gratton, 2018; Gyurkovics et al., 2022). Younger and older adults did not differ in the early phase of stimulus processing (Shift1), suggesting that they engage these early inhibitory mechanisms to a similar degree. Notably, the cue-induced spectral shift in the late processing phase (Shift3) was *still negative* (albeit to a lesser extent) in younger adults, suggesting that the momentary inhibition was followed by disinhibition (return to baseline) in this group, which may reflect their need to prepare to shift attention to the upcoming target. Conversely, in older adults, this later change was *positive*, suggesting an increased excitation following the early phase of inhibition. Shift1 showed broad scalp distribution in both age groups. In contrast, Shift3 showed a more fronto-central distribution in older adults (**Fig. 4C**), suggesting that the age-related excitation in the late processing phase involves changes in cortical activity that are more local and can be captured only at fronto-central sites in scalp-recorded EEG. A series of regression analyses shed further light on the mechanisms through which aperiodic activity is related to aging, stimulus processing, and behavior. Cue-induced spectral shifts predicted upcoming performance, with a more clockwise shift related to higher overall performance (as indexed by IES) and more efficient conflict resolution (as indexed by the congruency effect). As such, the regressions converge with the ANOVA findings, further supporting the interpretation of aperiodic activity as a viable marker of information processing that substantially contributes to subsequent behavior. Importantly, the strength of the relationship between the cue-induced spectral shifts and overall performance decreased when age was included in the models (**Fig. 5C**). This indicates that event-related spectral shifts can be considered a mixture of individual differences related to stimulus processing and aging, which additively shape overall performance (cf. Voytek et al., 2015).

### 4.3. Theoretical and Methodological Implications

The novel properties of the aperiodic background EEG reported here have important theoretical and methodological implications. First and foremost, the present results contribute to current theories of age-related cognitive decline (for reviews, see Fabiani et al., 2022; Grady, 2012; Jiang et al., 2023). In particular, the neural noise hypothesis of aging (Cremer & Zeef, 1987; Salthouse & Lichty, 1985; Voytek & Knight, 2015) posits that disrupted neural communication with advancing age and related inhibitory deficits – indexed by greater E:I ratio – become more pronounced after stimulus presentation, thereby reducing older adults’ ability to maintain newly formed representations. While several studies have attempted to address this hypothesis (Dave et al., 2018; Ribeiro & Castelo-Branco, 2022; Tran et al., 2020; Voytek et al., 2015), none are conclusive as they have not examined event-related changes in aperiodic activity, which greatly limits their interpretation in terms of information processing. In this study, despite replicating an age-related increase in the E:I ratio at baseline (the pre-cue window), older adults did not show significantly greater E:I ratio in the early phase of stimulus processing compared to younger adults (no age-group difference for Shift1). This indicates that there is no apparent deficit in the initial inhibitory response in older individuals, thus suggesting that the mechanism of age-related cognitive decline speculated so far may require some revision. Based on the observed age-related temporal changes in aperiodic activity and their relationships with performance, we propose that the greater *post-inhibitory excitation* observed in the late phase of information processing in older adults (an increased E:I ratio for Shift3) may be an excessive (i.e., greater than baseline) rebound after inhibition. This is consistent with the E:I framework (Gao et al., 2017; Gyurkovics et al., 2022; Waschke et al., 2021) and sheds new light on the origins of neural noise associated with stimulus processing (Voytek et al., 2015; Voytek & Knight, 2015).

Relatedly, the ANOVA results also suggest that the age-related increase in the E:I ratio can be experimentally counteracted by providing older adults with greater contextual diversity and novelty (frequently changing cues in this study), which triggers a heightened level of performance. It will be important for future research to test how long event-induced aperiodic changes persist and what other forms of experimental manipulations can help overcome the age-related E:I imbalance towards excitation. As here we focused on cue-induced (proactive) processes and the target-locked EEG was deliberately excluded (as it was contaminated with manual responses), future research would also benefit from tracking the dynamics of aperiodic activity in response to an imperative stimulus, provided that contamination from motor activity can be excluded.

At the methodological level, this study reinforces the notion that the ERPs contribute to the broadband EEG background activity (Gyurkovics et al., 2022), emphasizing the need for their removal before estimating the 1/*f*^x^ (aperiodic) parameters. Furthermore, our findings greatly extend the current understanding of event-related shifts in aperiodic activity by revealing their temporal variability and offering a viable methodological framework for studying dynamic changes in the E:I balance over time. Although the ideal length of the time window for quantifying spectrograms is still an open research question, we demonstrated that a 500-ms temporal integration window provides a robust and effective method for quantifying temporal aperiodic changes in scalp EEG. Moreover, to ensure the highest data quality, we employed rigorous EEG quality control, including careful assessment of spectrograms and spectral parameterization outcomes. We also utilized a relatively large sample size (compared to typical studies in this field), which further increases the statistical power of the analyses and improves the generalizability of our findings. Collectively, the results presented here indicate that a 500-ms temporal integration window, along with strict data quality control and a relatively large sample size, offer a robust and effective framework for quantifying temporal aperiodic changes in the scalp EEG recordings, thus providing a promising avenue to better understand the brain dynamics underlying information processing.

Finally, the evidence for dynamic changes in the aperiodic component reconciles seemingly conflicting reports regarding attention-dependent spectral changes in scalp-recorded EEG. While Gyurkovics et al. (2022) reported an attention-dependent exponent *increase,* Waschke et al. (2021) reported an attention-dependent exponent *decrease*. One of the methodological differences between these studies is that they focused on the aperiodic activity from different post-stimulus periods. Gyurkovics and colleagues focused on the immediate response to the stimulus, whereas Waschke and colleagues quantified the spectrum several hundred milliseconds after stimulus onset and did not control for any lingering ERP contributions. The current results suggest that the discrepancy between these two previous studies may only be coincidental. Here, compared to the pre-event period, the exponent was more negative immediately after the stimulus, consistent with Gyurkovics et al., and less negative in the furthest time window, consistent with Waschke et al. (**Fig. 3A**). Given this apparent discrepancy and the risk of misinterpretation, future studies should account for the dynamic nature of the aperiodic activity or at least carefully address the period over which they quantify the spectra. This seems all the more important given that the regression analyses showed that the exponent can convey different information depending on the time window in which it is quantified. Although more research is needed on this topic, aperiodic neural activity immediately after the stimulus seems to be the most sensitive to experimental effects, whereas later activity may also reflect the contribution of individual differences, such as those due to aging.

### 4.4. Conclusions

To our knowledge, this study is the first to investigate the temporal dynamics of broadband (aperiodic) EEG background activity during stimulus processing in younger and older adults. Our findings show that the spectral exponent – an overarching measure of the shape of the broadband EEG – is not a stationary feature of electrophysiological signals but a dynamically changing phenomenon that provides insights into the neural bases of stimulus processing and its changes with aging. From a theoretical standpoint, these data contribute to neuroscientific models of cognitive processing and age-related cognitive decline. From a methodological standpoint, the study provides a viable framework for investigating the temporal dynamics of aperiodic activity and the alternation of excitation and inhibition in neural circuits, providing cross-scale links with single and multiple-unit activity and imaging research.

## Acknowledgments

This work was supported by NIA grant RF1AG062666 to G. Gratton and M. Fabiani. P. Kałamała was supported by National Science Centre Poland grant 2020/36/T/HS6/00363. We acknowledge Brooke Frazier, Dana Joulani, Rebecca Lii, Madeleine Peckus, Preeti Subramaniyan, and Yunsu Yu for their help with data collection.

## Footnotes

1 The pattern of results replicates when incorrect trials are excluded.

